# A quick and cost-effective method for DNA-free total RNA isolation using magnetic silica beads

**DOI:** 10.1101/2020.11.04.341099

**Authors:** Aniruddha Das, Debojyoti Das, Arundhati Das, Amaresh C. Panda

## Abstract

Current RNA purification methods widely use silica-based columns that allow quick isolation of high-quality and good quantities of RNA. However, the major limitations include high cost, the requirement of different kits for small RNA isolation, genomic DNA contamination, and not being flexible. Here, we used the in-house RNA isolation reagent (RIR) for cell lysis, followed by RNA precipitation using isopropanol. RNA isolated using the in-house RIR resulted in a similar quantity and quality compared to the commercial TRIzol. Furthermore, the commercial RNA isolation kits with silica-based columns recommend genomic DNA digestion during or after RNA purification, adding time and cost to RNA purification. Here, we developed an optimized in-house protocol for isolating high-quality RNA free of genomic DNA contamination using magnetic silica beads without needing DNase digestion. Additionally, our method purifies total RNA along with the small RNA fraction, including miRNAs, which usually require a separate kit for extraction. Additionally, the RNA prepared with our method was equally suitable for mRNA and miRNA expression analysis using RT-qPCR. Together, the in-house method of RNA isolation using the magnetic silica beads has exhibited comparable or better total RNA extraction compared to commercial kits at a fraction of the cost and across various cells and tissues.

## INTRODUCTION

The quality and quantity of RNA are the first and most crucial steps for ensuring downstream gene expression analysis accurately. The integrity of the total RNA is checked before its use in downstream applications, including quantitative RT-PCR, microarrays, and RNA sequencing, which are expensive, time-consuming, and labor-intensive (Tan & Yiap, 2009). The acceptable total RNA for downstream applications must fulfill a few requirements, including the RNA preparation must be free from protein, salts, alcohol, phenol, and genomic DNA (gDNA), which may interfere with downstream enzymatic reactions and gene expression analysis (Bustin & Nolan, 2004; Schrader, Schielke, Ellerbroek, & Johne, 2012; Tan & Yiap, 2009; Wilson, 1997). Additionally, the total RNA must be free of nucleases and stored appropriately to maintain integrity.

Various RNA extraction protocols have been developed to isolate high-quality RNA from various cells and tissues. The RNA extraction methods broadly fall into three categories: organic extraction followed by RNA precipitation, purification through a silica-membrane column, and silica-coated magnetic particles. Although the classical acid guanidinium thiocyanate-phenol-chloroform (AGPC) method of RNA extraction followed by RNA precipitation has been the most commonly used method for decades, the isolated RNA samples are often contaminated with proteins, gDNA, organic solvents, and salts (Chomczynski & Sacchi, 2006; Rio, Ares, Hannon, & Nilsen, 2010b; Schrader et al., 2012; Tan & Yiap, 2009; Wilson, 1997). However, the popular commercial kits use guanidinium salt-based lysis buffers without toxic organic solvents, followed by RNA purification with silica columns and magnetic silica beads (Rio, Ares, Hannon, & Nilsen, 2010a; Yaffe et al., 2012). These commercial kits using the silica column are effective, relatively simple, and yield sufficient quantities of intact RNA. However, the silica columns often lose the small RNA fractions and recommend using specific RNA isolation kits to isolate miRNAs. The magnetic bead-based RNA purification method is relatively new, and the kits are mostly available for specific applications such as viral RNA isolation (Berensmeier, 2006; P. Oberacker et al., 2019). Additionally, magnetic bead-based kits are often supplied with an automatic high-throughput RNA isolation system, which is very expensive. Additionally, the silica column and magnetic bead-based purification methods often result in gDNA contamination in the purified RNA (Bustin & Nolan, 2004; Rio, Ares, Hannon, & Nilsen, 2010c; Tan & Yiap, 2009). Since all these RNA isolation methods do not remove DNA, DNase treatment is recommended to remove the contaminating gDNA before proceeding to downstream applications, which adds significant time and cost to RNA purification (Rio et al., 2010c).

In this work, we demonstrated a method where cells are lysed with the in-house prepared AGPC-based cell lysis followed by depletion of the gDNA using a silica spin column. After gDNA depletion, the cell lysate was mixed with ethanol and allowed to bind silica-coated magnetic beads, followed by washing and eluting DNA-free total RNA. This method also isolates the small RNAs, including miRNAs, for expression analysis. Together, the proposed method is quick, practical, and cost-effective for isolating DNA-free total RNA, including small RNAs.

## MATERIALS AND METHODS

### HeLa, C2C12, and βTC6 cell culture

Mouse C2C12, βTC6, and human HeLa cells were cultured in DMEM supplemented with 10/15% FBS and antibiotics. The cells were maintained in a 5% CO2 humidified atmosphere at 37°C. The cells were washed with 1X PBS and trypsinized with 0.25% Trypsin-EDTA, then pelleting the desired number of cells for RNA isolation.

### RNA isolation reagents and kits

The commercial RNA isolation kits were procured from different companies, including TRIzol (Thermo Fisher Scientific, 15596026), miRNeasy Mini Kit (Qiagen, 217004), PureLink miRNA isolation kit (Thermo Fisher Scientific, K157001), and HiPurA™ Total RNA Miniprep Purification Kit (HiMedia, MB602-250PR). The on-column DNase I digestion set (Sigma, DNASE70) was used for genomic DNA (gDNA) digestion during RNA purification. The gDNA depletion silica columns (EconoSpin silica columns, 1920) were procured from Epoch life sciences. The MagPrep Silica Particles (Sigma-Aldrich, 101193) were purchased from Sigma for RNA isolation. The in-house RNA isolation reagent (RIR) comprises 38 % phenol, 0.4 M ammonium thiocyanate, 0.8 M guanidine thiocyanate, 0.1 M sodium acetate, and 5 % glycerol. RNA wash buffer comprised 10 mM Tris-HCl pH 7.5 and 75% ethanol. All reagents for buffer preparation were procured from Sigma, SRL, or HiMedia.

### RNA isolation

The first method is the classical AGPC-based RNA isolation method described previously (Chomczynski, 1993; Chomczynski & Sacchi, 2006). Briefly, HeLa cells were lysed in 500 μl TRIzol (Thermo Scientific) or in-house RIR by pipetting, followed by mixing with 100 μl of chloroform. Next, the mixture was centrifuged at 12,000x g for 10 min at 4 °C, followed by a collection of 200 μl of the aqueous layer in a new tube. Next, the aqueous layer was mixed with 200 μl of isopropanol and incubated for 10 min at room temperature, followed by centrifugation at 12,000x g for 10 min at 4 °C. Then, the supernatant was discarded, and the pellet was washed with the wash buffer (75% ethanol) and centrifuged at 12,000x g for 5 min at RT. Finally, the pellet was air-dried for 3-5 min and dissolved in 50 μl nuclease-free water.

In the in-house method, two to five million cells were lysed in 500 μl RIR by vigorous pipetting followed by mixing with 100 μl of chloroform and centrifuged at 12,000x g for 10 min at 4 °C. The 200 μl aqueous layer was transferred into a new tube. The flowthrough was passed through a silica column by centrifugation at 12,000x g for 1 min at room temperature. Next, the flowthrough was mixed with 300 μl of ethanol and 10 μl of MagPrep silica beads. The mixture was incubated for 5 min at room temperature with shaking at 1200 rpm on a thermomixer, followed by putting the tube on a magnetic stand, separating the beads to one side of the tube, and the supernatant was discarded. For DNase treatment, the beads were digested for 15 min with DNase I following the manufacturer’s protocol. Next, the beads were washed twice with 500 μl wash buffer. The wash buffer was discarded by holding the beads on a magnetic stand, followed by the repetition of this step after a short spin. The tube was incubated at room temperature for about 3 minutes to allow the ethanol to evaporate entirely. The beads were then mixed with 40 μl nuclease-free water and incubated for 2-3 minutes before putting the tube on the magnetic stand. The dissolved RNA was collected in a fresh tube for quality and quantity assessment.

Also, we used the HiPurA Total RNA Miniprep Purification Kit (cat no. MB602-250PR; HiMedia), miRNeasy Mini Kit (cat no. 217004; Qiagen), and PureLink miRNA isolation kit (cat no. K157001; Thermo Fisher) for preparing the total RNA and miRNA following the manufacturer’s instructions.

### Assessment of RNA quantity and quality

The quantity and quality of RNA samples were analyzed using NanoDrop 2000 or Multiskan Sky (Thermo Scientific) by determining the A260/A280 and A260/A230 ratios (Manchester, 1996). The gDNA contamination in RNA was also assessed with the Qubit 4 using a dsDNA BR assay kit. The RNA integrity and gDNA contamination were evaluated by resolving the RNA on an SYBR-Gold stained 1.2 % denaturing formaldehyde-agarose gel and visualizing it under a UV transilluminator. Furthermore, RNA samples were analyzed on a TapeStation 4200 (Agilent) using an RNA ScreenTape to determine the RNA integrity number (RIN).

### Reverse transcription followed by quantitative PCR (RT-qPCR) and agarose gel analysis

For long RNA analysis, 1 μg of total RNA was isolated using different kits, and the in-house MagPrep silica beads method was used to synthesize cDNA using the High Capacity cDNA Reverse Transcription (RT) Kit (Thermo Fisher Scientific) following the manufacturer’s protocol. The cDNA was diluted to 500 μl with nuclease-free water, and 2X PowerUP SYBR Green Master Mix (Applied Biosystems) was used for RT, followed by quantitative PCR (RT-qPCR) analysis using convergent primers for linear and divergent primers for circular RNAs (**Supplementary Table S1**) (Das, Das, & Panda, 2022). For analyzing the small RNA fractions, total RNA was used for cDNA synthesis with the Mir-X miRNA First-Strand Synthesis Kit (Clontech) following the manufacturer’s protocol. Since the PureLink miRNA isolation kit isolates the long RNAs in column 1 and small RNAs (miRNA) in column 2, we mixed an equal volume of both RNAs and considered that as total RNA for miRNA cDNA synthesis. The miRNA cDNA samples were diluted to contain cDNA from 0.5 ng of RNA/μl. The small RNA levels were analyzed by RT-qPCR analysis using small RNA-specific primers (**Supplementary Table S1**) and the primers provided in the Mir-X kit. RT-qPCR amplification was performed at 95 °C for 2 minutes and subsequently 40 cycles of 95 °C for 2 s and 60 °C for 20 s on a QuantStudio Real-Time PCR System (Thermo Fisher Scientific). The relative expression of different target RNAs in RNA samples isolated with different methods was analyzed using their Ct values in RT-qPCR. RT-PCR amplification was performed using the 2X DreamTaq PCR mastermix (Thermo Fisher Scientific) with a cycle set up of an initial 95 °C for 2 minutes, followed by 40 cycles of 95 °C for 2 s and 60 °C for 20 s on a thermal cycler. The amplified PCR products were resolved in an SYBR-Gold stained 2.5 % agarose gel and visualized by a UV transilluminator gel documentation system.

## RESULTS AND DISCUSSION

### RNA preparation with in-house RIR yields comparable quality and quantity of RNA compared to TRIzol

Here, we wanted to analyze the quantity and quality of RNA extracted using the classical TRIzol extraction method and the in-house RIR (Rio et al., 2010b; Walker & Lorsch, 2013). Therefore, we used three cell lines, including HeLa, βTC6, and C2C12 cells, to isolate total RNA using in-house RIR and TRIzol with the standard RNA isolation protocol using isopropanol precipitation of the aqueous layer. Each replicate was divided into equal halves to prepare RNA simultaneously with TRIzol and RIR. As expected, the quality and quantity obtained with the in-house RIR were comparable with the commercial TRIzol (**Figure 1**).

**Figure 1.**
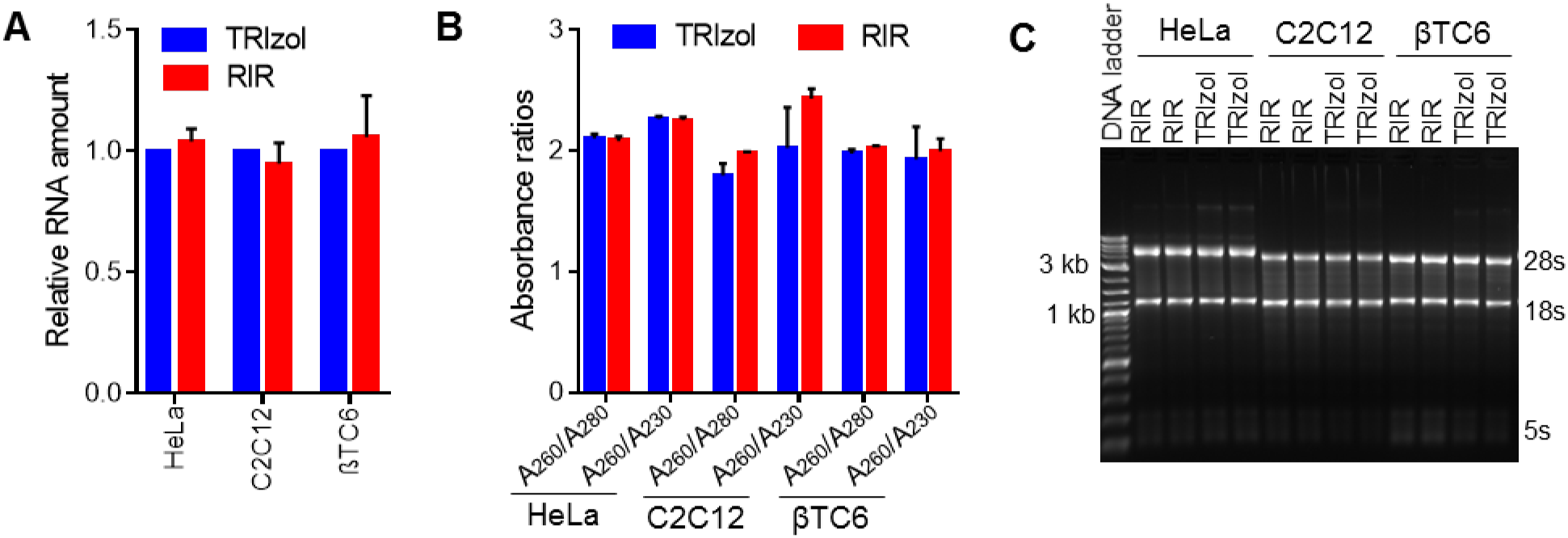
Comparison of RNA isolated using RIR and TRIzol. **A**. The relative amount of total RNA recovered using RIR and TRIzol from HeLa, C2C12, and βTC6 cells. **B**. The purity of total RNA prepared using RIR and TRIzol from mentioned cell lines was measured by the ratios of A260/A280 and A260/A230. **C**. Total RNA was prepared with RIR and TRIzol from mentioned cells and was resolved on a denaturing 1.2% agarose gel stained with SYBR-Gold. Data in panels A and B represent the mean ± SEM from 6 independent experiments.

Next, we wanted to use the aqueous layer from RIR to prepare RNA using magnetic silica beads. As shown in **Figure 2A**, the aqueous layer was incubated with MagPrep silica magnetic beads to adsorb the RNA, followed by digestion with on-column DNase I. Finally, the DNase-digested beads were washed, and the bound RNA was eluted from the beads. HeLa and C2C12 cells were used for RNA extraction using MagPrep silica beads. However, DNase I treatment reduced the total amount of RNA extracted from HeLa and C2C12 cells (**Figure 2B**).

**Figure 2.**
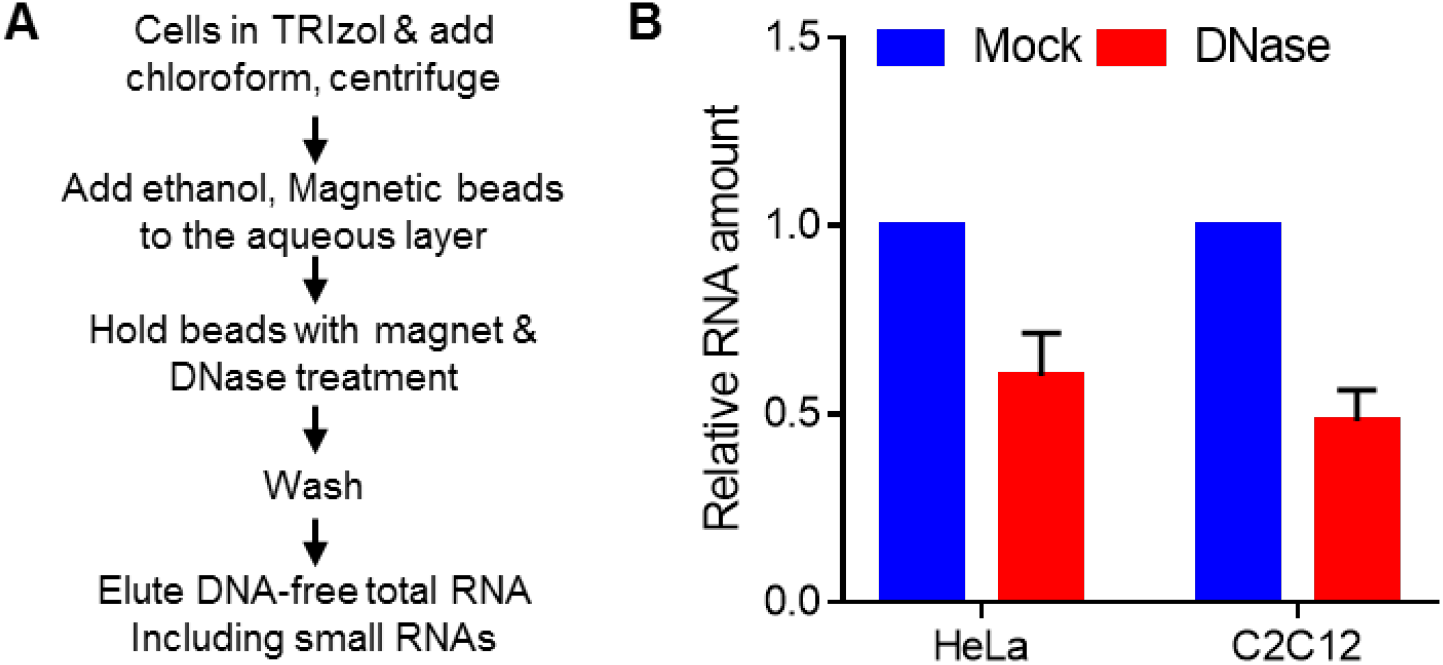
DNase digestion during the RNA preparation with MagPrep silica beads decreases the RNA yield. **A**. Flow chart of crucial steps of in-house optimized RNA extraction method using magnetic silica beads. **B**. Amount of total RNA recovered with or without DNase digestion from HeLa and C2C12 cells using an in-house optimized MagPrep silica beads RNA extraction protocol, as shown in panel A. Data represent the mean ± SEM from 3 independent experiments.

### Removal of genomic DNA with silica column eliminates the time-consuming and costly DNase digestion step

As shown in **Figure 3A**, we passed the aqueous layer from the RIR-chloroform step through a silica column to remove the genomic DNA, followed by RNA purification using the MagPrep silica magnetic beads. Unlike the reduction in RNA yield after DNase treatment, gDNA depletion using the silica column did not affect the RNA yield (**Supplementary Figure S1A**). Also, gDNA depletion completely removed the gDNA contamination from total RNA isolated using MagPrep silica beads from HeLa and C2C12 cells (**Supplementary Figure S1B**). We found that cell lysis with RIR, followed by gDNA removal using the silica column, quickly and inexpensively removes gDNA without needing expensive and time-consuming DNase digestion. Isolation of DNA-free total RNA using the gDNA depletion column followed by RNA isolation with MagPrep silica beads could isolate more than 20 μg of total RNA from 5 million HeLa cells (**Figure 3B**). The isolation of DNA-free total RNA from HeLa, C2C12, and βTC6 cells showed clean 28s, 18s, and 5s rRNAs on the gel, suggesting the isolation of good-quality RNA for downstream analysis (**Figure 3C)**.

**Figure 3.**
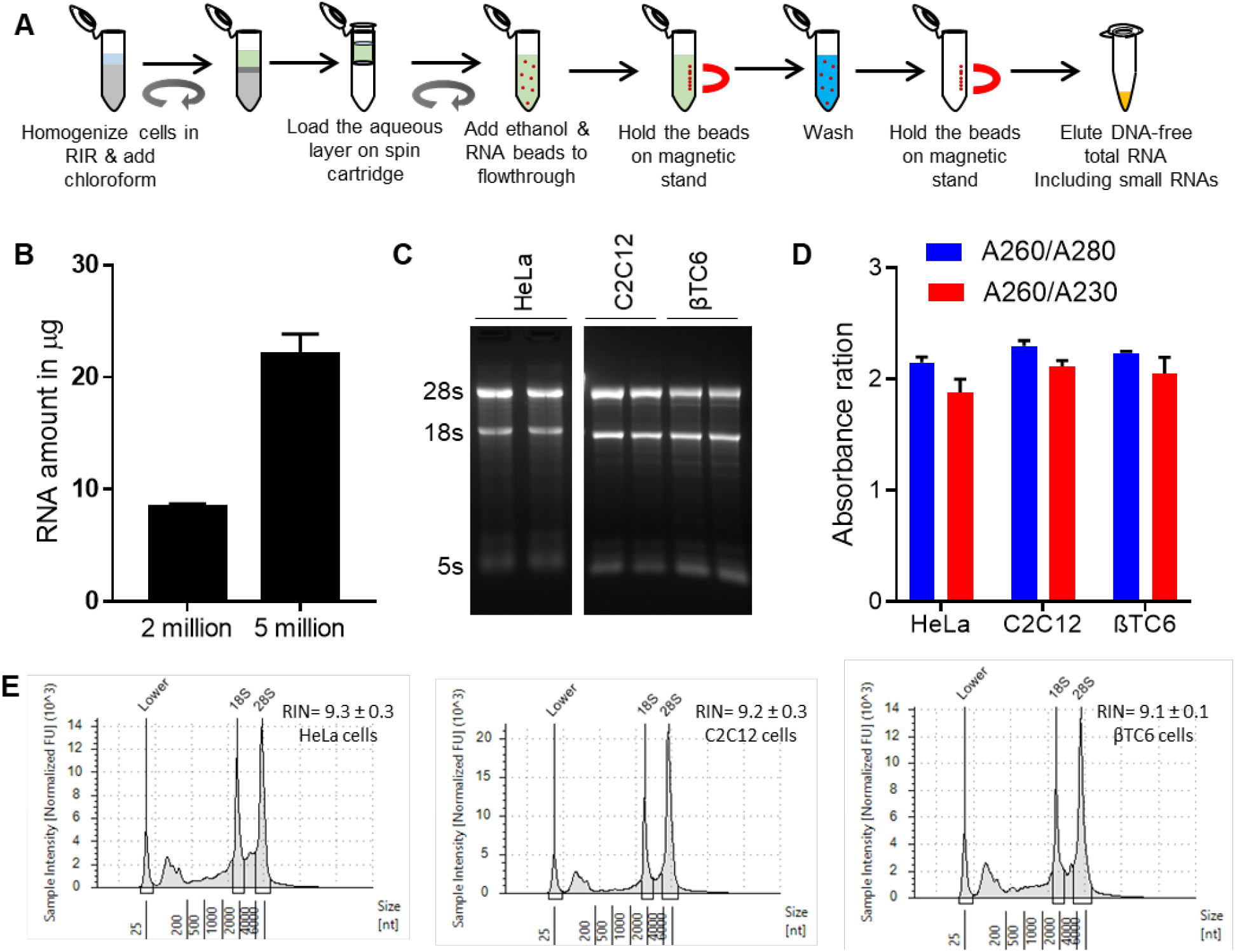
RNA preparation with magnetic silica beads. **A**. Flow chart of crucial steps of in-house optimized DNA-free total RNA extraction method using MagPrep silica beads. **B**. DNA-free total RNA recovered using MagPrep silica beads from 2 and 5 million HeLa cells. **C**. Denaturing 1.2% agarose gel depicting the total RNA from HeLa, C2C12, and βTC6 cells prepared using MagPrep silica RNA beads with or without gDNA depletion. **D**. The ratio of A_260_/A_280_ and A_260_/A_230_ of total RNA recovered using MagPrep silica beads from HeLa, C2C12, and βTC6 cells. **E**. Tapestation analysis showing representative electropherogram of DNA-free total RNA extracted from HeLa, C2C12, and βTC6 cells using MagPrep silica beads. Data in panels B, D, and E represent the mean ± SEM from at least 3 independent experiments.

Furthermore, the purity of DNA-free total RNA prepared with the in-house method using MagPrep silica beads was analyzed by calculating the ratio between the absorbance at 260 nm vs. 280 nm (A260/A280) and 260 nm vs. 230 nm (A260/A230). As shown in **Figure 3D**, all the 260/280 ratios for RNA prepared using magnetic silica beads from HeLa, C2C12, and βTC6 cells were around 2.0, suggesting that the isolated RNA was without DNA or protein contaminants. Additionally, the 260/230 ratios were between 2-2.3, indicating high-quality RNA with no contaminants that can be used in downstream applications (Manchester, 1996). Although the spectroscopy method can determine the quality and quantity of RNA, it cannot determine the integrity of the RNA. Our TapeStation analysis of RNA integrity showed little to no sign of degradation with RIN values of more than 9 for total RNAs isolated from HeLa and C2C12 cells (**Figure 3E**). Total RNAs with RIN values of more than 6.4 are recommended for transcriptome analysis using RNA sequencing (Gallego Romero, Pai, Tung, & Gilad, 2014). Therefore, the DNA-free total RNA prepared with the MagPrep beads is suitable for downstream analysis, including reverse transcription, RT-PCR, and RNA sequencing.

### RNA prepared with MagPrep silica beads is equally good for gene expression analysis compared to commercial RNA isolation kits

To compare the functionality and suitability of the RNAs extracted from C2C12 cells using our MagPrep silica beads, we prepared good quality RNAs using the TRIzol, miRNeasy Mini Kit, PureLink miRNA isolation kit, and HiPurA Total RNA Miniprep Purification Kit (**Figure 4A**). Additionally, the gel image indicated that RNA purified with our magnetic bead-based method successfully isolated the small RNAs, which was hardly visible in some of the RNA prepared with the commercial kit (**Figure 4A)**. Furthermore, the total RNAs were reverse transcribed, and specific mRNA and circRNAs were successfully amplified, suggesting that the RNA prepared with the in-house MagPrep RNA isolation method is as good as the commercial kit for RT-PCR analysis (**Figure 4B**). Furthermore, RT-qPCR analysis of *Gapdh* mRNAs, *circSamd4, circRras2*, and *circTcf20* showed similar expression levels in RNA prepared with our MagPrep method compared to the commercial kits (**Figure 4C)**. Since the PureLink miRNA isolation kit prepares long and small RNA separately, we mixed an equal volume of long and small RNA to obtain the total RNA for miRNA analysis. RT-qPCR analysis showed that the RNA prepared with the MagPrep method isolates miRNAs with similar or better efficiency than the commercial RNA isolation kits (**Figure 4D)**. Together, total RNA preparation with the in-house method using magnetic silica beads could be used for the expression analysis of all types of RNAs, such as mRNAs, circRNAs, and miRNAs.

**Figure 4.**
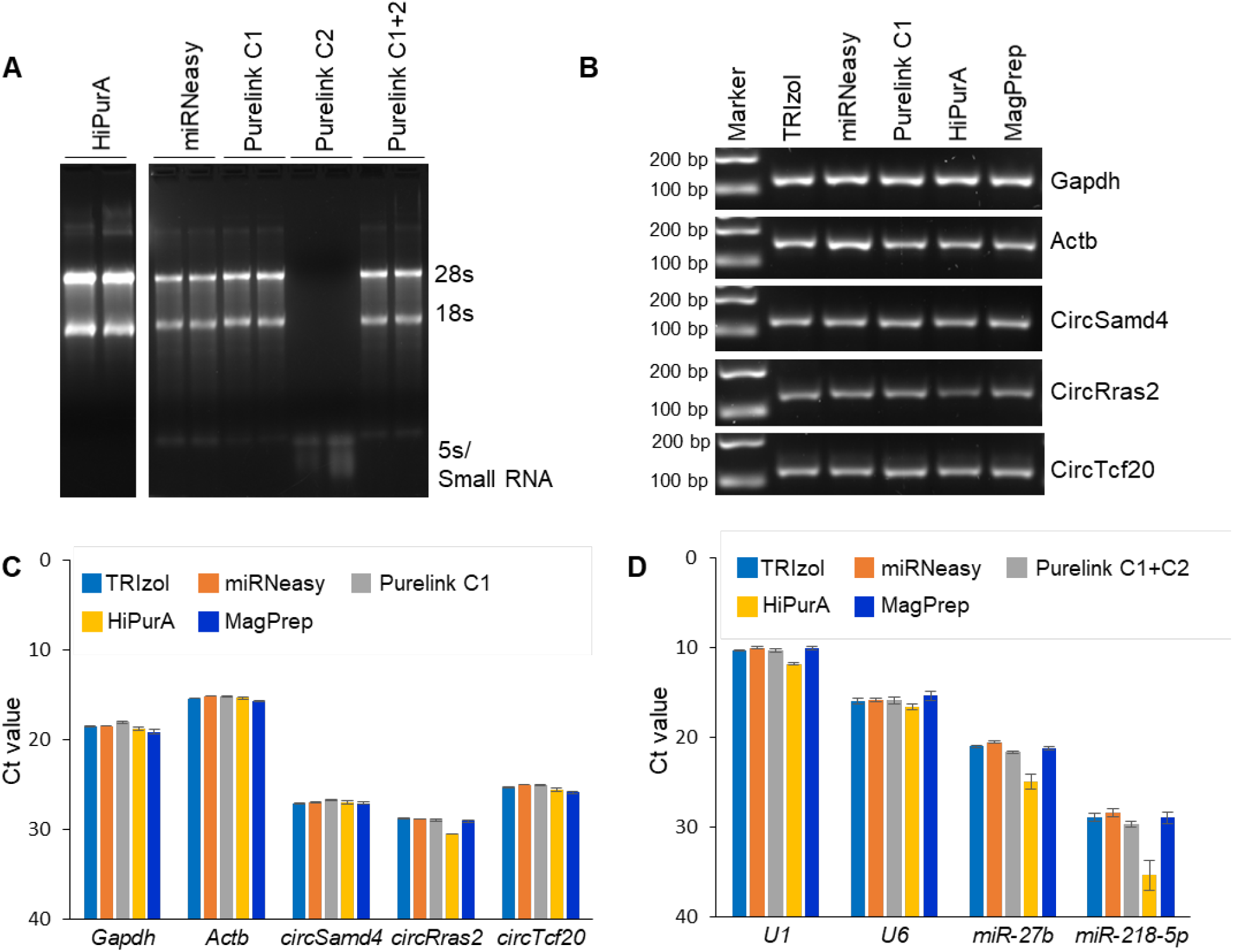
RNA prepared with the in-house MagPrep method is equally good compared to commercial RNA isolation kits for gene expression analysis. **A**. Denaturing 1.2% agarose gel depicting the total RNA or miRNA prepared with different commercial kits using silica columns. Purelink C1 represents long RNA isolated from column 1, Purelink C2 represents small RNA isolated from column 2, and C1+C2 represents the mixture of RNA isolated from column 1 and column 2 of the Purelink kit. **B**. RT-PCR products of *Gapdh, Actb, circSamd4, circRras2*, and *circTcf20* were visualized in an SYBR Gold stained 2.5 % agarose gel. **C**. RT-qPCR analysis and Ct values of mRNAs and circular RNAs including, *Gapdh, Actb, circSamd4, circRras2*, and *circTcf20* in 10 ng of total RNA prepared with different commercial kits and in-house MagPrep method. **D**. RT-qPCR analysis and Ct values of small RNAs, including *U1* snRNA, *U6* snRNA, *miR-27b*, and *miR-218-5p* in 2.5 ng of total RNA prepared with different commercial kits and in-house MagPrep method. Data in panels C and D represent the means ± SEM from three independent experiments.

## CONCLUSIONS

The isolation of intact and pure RNA is critical for all downstream applications, including cDNA synthesis, PCR, real-time PCR, northern blotting, microarray, and RNA sequencing (Tan & Yiap, 2009). There are three commonly used methods of RNA purification, such as acid-phenol extraction, silica column, and magnetic silica bead-based isolation. The classical organic extraction of RNA using TRIzol is the cost-effective and straightforward gold standard method for RNA isolation (Rio et al., 2010b). However, it has a few disadvantages, including using hazardous reagents, time-consuming, laborious protocol, and often contamination with salts and other biomolecules (Bustin & Nolan, 2004; Wilson, 1997). On the other hand, the silica-based RNA purification column is the most commonly used in commercial kits, isolating high-quality and right amounts of RNA without using the toxic-reagents (Rio et al., 2010a). Consequently, RNA isolation with magnetic silica beads is available for specific applications, including viral RNA isolation and automatic high-throughput applications (P. Oberacker et al., 2019). In addition, the magnetic silica bead-based high-throughput RNA isolation kits require expensive instruments and consumables to automate the process.

Furthermore, the major limitation of the silica column or magnetic bead-based commercial RNA isolation kits is the incomplete lysis, contamination of RNA with gDNA, and high cost (Bustin & Nolan, 2004; Wilson, 1997). Since the downstream gene expression analysis application strictly demands DNA-free RNA, some commercial kits include the on-column DNase digestion step, and others recommend DNase treatment after RNA purification. However, DNase treatment adds significant time and cost to the experiment. In addition, DNase treatment often leads to a loss of RNA amount and might affect RNA integrity due to high-temperature exposure during heat inactivation of commercial DNases. Furthermore, the silica columns in commercial kits often miss the small RNA fractions, and specific kits are required to isolate miRNAs (Rio et al., 2010a). To overcome all these shortcomings with the commercial kits, we developed a quick, efficient, flexible, and inexpensive RNA extraction method. The user can optimize the amount of TRIzol, RIR, or commercial RNA lysis buffer depending on the input sample type and amount. First, the aqueous phase is passed through the gDNA depletion silica column removing the gDNA without using a time-consuming and expensive DNase treatment procedure (**Figure 3**). Then, the lysate was mixed with ethanol and MagPrep silica beads to bind the RNA in the sample. The user can choose the amount of beads depending on the expected amount of RNA in the sample. The proposed method used magnetic beads for RNA isolation with only one washing step, reducing the hands-on time and simplifying the protocol. The commercial MagPrep silica beads can be replaced with in-house prepared silica-coated magnetic beads (Phil Oberacker et al., 2019).

Furthermore, RNA purity analysis indicated that the RNA prepared with this method is free of gDNA, salt, and protein contaminant (**Figure 3**). The analysis of RNA integrity on gel and TapeStation indicated the isolation of high-quality, intact RNAs for downstream applications. The RT, followed by PCR, shows that the RNA prepared with our proposed method is compatible with downstream applications like reverse transcription and PCR (**Figure 4**). Interestingly, RT-qPCR analysis suggested similar or better recovery and amplification of smaller-size RNAs with our MagPrep RNA than some commercial column-based kits. Furthermore, RNA samples isolated through our method could efficiently isolate small RNA fractions for miRNA analysis, which often requires a separate kit. Our RNA isolation procedure can be completed in about 30 min, which is significantly less than the commercial kits to get DNA-free RNA, including miRNAs. In summary, we believe that the MagPrep RNA isolation method is a quick, effective, and cost-effective in-house method for preparing DNA-free total RNA compared to the existing commercial kits.

## Acknowledgments

The authors thank our colleagues at the Institute of Life Sciences, Bhubaneswar, for helpful discussions and suggestions to improve the method.

## Authors Contribution

Conceptualization: Amaresh C. Panda; Methodology: Aniruddha Das, Debojyoti Das, Arundhati Das; Formal analysis and investigation: Aniruddha Das, Debojyoti Das, Arundhati Das; Writing - original draft preparation: Aniruddha Das, Debojyoti Das, Arundhati Das; Writing - review and editing: Aniruddha Das, Debojyoti Das, Arundhati Das, Amaresh C. Panda; Funding acquisition: Amaresh C. Panda; Resources: Amaresh C. Panda; Supervision: Amaresh C. Panda.

