## Supplementary Figure S1 for "A quick and cost-effective method for DNA-free total RNA isolation using magnetic silica beads"

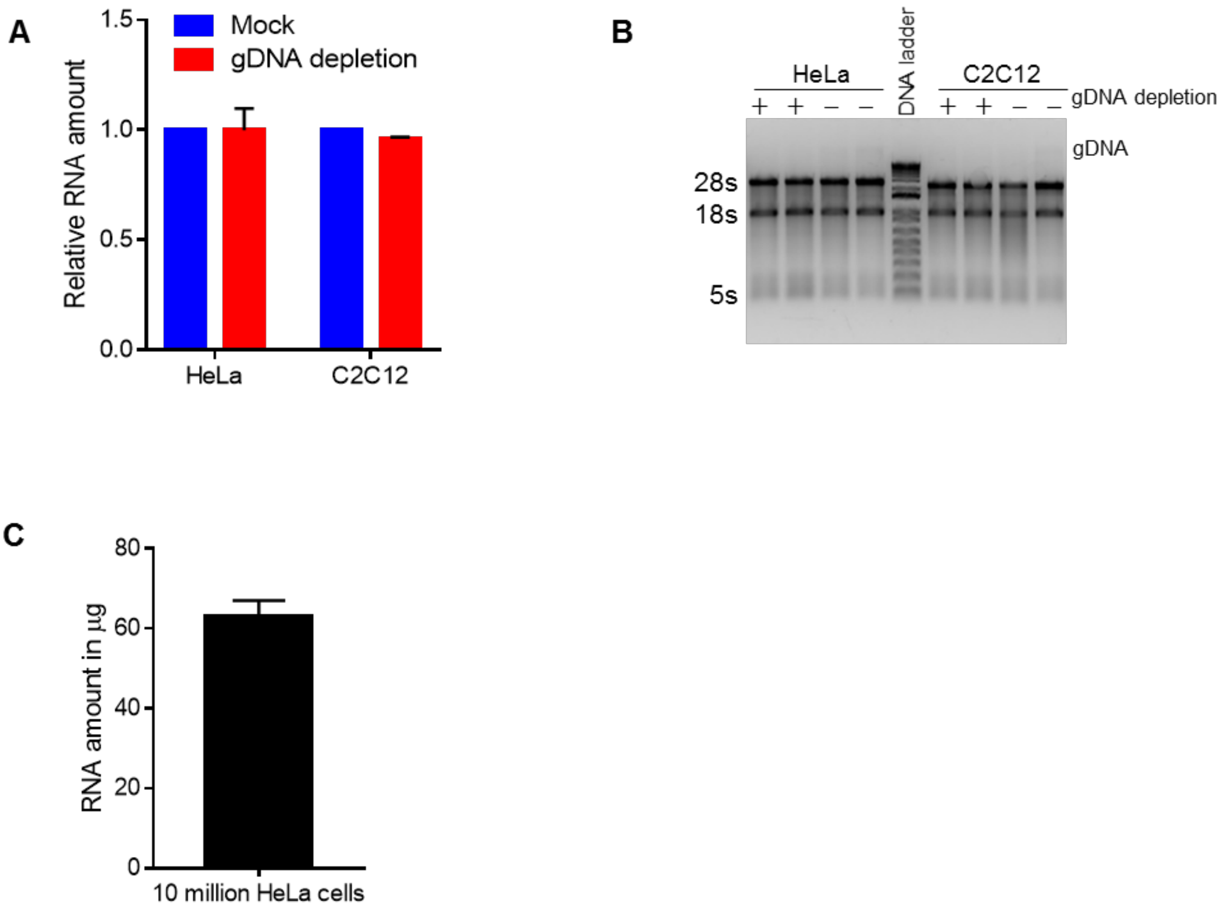


**Supplementary Figure S1. RNA preparation with magnetic silica beads requires gDNA depletion with DNase or silica column. A.** Amount of total RNA recovered with or without using the gDNA depletion column. The aqueous layer from the RIR-chloroform step was either passed through the gDNA depletion column or was used directly for RNA extraction using MagPrep silica beads. **B.** Total RNA prepared using MagPrep silica beads with or without using the gDNA depletion column were resolved on a denaturing 1.2% agarose gel stained with SYBR-Gold. **C.** Total RNA was recovered using MagPrep silica beads from 10 million HeLa cells with gDNA depletion. Data in panels A and C represent the mean ± SEM from 3-5 independent experiments.
