## Supplementary Table S1 for "A quick and cost-effective method for DNA-free total RNA isolation using magnetic silica beads"

**Supplementary Table S1.** Oligonucleotides used in the study

| Primer name | Sequence (5' to 3') |
|---|---|
| Gapdh-F | AACTTTGGCATTGTGGAAGG |
| Gapdh-R | GGATGCAGGGATGATGTTCT |
| Actb-F | TGTTACCAACTGGGACGACA |
| Actb-R | GGGGTGTTGAAGGTCTCAAA |
| circSamd4-F | GCCCCTCTACTCTTCCTCGT |
| circSamd4-R | CAAAGGCAGGTGAGTTAGCA |
| circRras2-F | AAGGACCGTGATGAGTTTCC |
| circRras2-R | GGTCATCGATCACACATTGC |
| circTcf20-F | ACCATTACCCATGTGCCATT |
| circTcf20-R | CTCCTGTGGGTAGCTCTGCT |
| U1-F | TACTTACCTGGCAGGGGAGATAC |
| U1-R | GAACGCAGTCCCCCACTAC |
| miR-27b | TTCACAGTGGCTAAGTTCTGC |
| miR-218-5p | TTGTGCTTGATCTAACCATGT |
